# MinION nanopore sequencing provides similar methylation estimates to Sanger bisulfite sequencing in the *TRPA1* promoter region

**DOI:** 10.1101/2021.09.17.460763

**Authors:** Sara Gombert, Kirsten Jahn, Hansi Pathak, Alexandra Burkert, Gunnar Schmidt, Lutz Wiehlmann, Colin Davenport, Björn Brändl, Franz-Josef Müller, Andreas Leffler, Maximilian Deest, Helge Frieling

**Author notes:** Sara Gombert and Kirsten Jahn contributed equally. Maximilian Deest and Helge Frieling contributed equally.

## Abstract

Bisulfite sequencing has long been considered the gold standard for measurement of DNA methylation at single CpG resolution. In the meantime, several new approaches have been developed, which are regarded as less error-prone. Since these errors were shown to be sequence-specific, we aimed to verify the methylation data of a particular region of the *TRPA1* promoter obtained from our previous studies. For this purpose, we compared methylation rates obtained via direct bisulfite sequencing and nanopore sequencing. Thus, we were able to confirm our previous findings to a large extent.

## 1. Introduction

Bisulfite sequencing, developed by Frommer and colleagues [1], is a method based on the sodium bisulfite-mediated conversion of cytosine to uracil in single-stranded DNA, and has long been considered the gold standard for methylation analysis. However, this method is susceptible for errors due to the bisulfite conversion as well as the subsequent amplification of DNA strands, which can lead to misinterpretation of the results. The harsh chemical treatment of DNA leads to significant degradation, thereby causing bisulfite conversion errors [2, 3]. Therefore, a balanced control between the desired conversion of unmethylated cytosines to uracils and the undesired DNA degradation and inappropriate conversion of methylated cytosines to thymines is indispensable. Otherwise, the unpredictable level of false positive and false negative results may be elevated due to the differing conversion efficiencies of cytosines depending on the sequence context [2]. However, when using modern kits for bisulfite treatment, conversion errors are relatively low [4, 5], whereas recovery rates are between 18 and 50 % [5]. The amplification of the target regions in bisulfite sequencing imposes the risk to intensify biases, especially due to the elevated error rates in high- and low-GC regions [6]. The amplification of artefacts from sequence-specific bisulfite-induced degradation and conversion errors leads to a higher overall bias in protocols involving amplification [7]. Since some regions are more susceptible to biases than others [7], the error-proneness of a sequence of interest is unpredictable. The choice of bisulfite conversion protocol or polymerase significantly reduces these artefacts but cannot completely abolish them [7]. The novel method of nanopore sequencing does not involve such error-prone procedures like bisulfite treatment or amplification of target regions. Nanopore sequencing was shown to discriminate between the four standard bases by measuring the change in current as the DNA /RNA molecule translocates through a protein nanopore. Methylated cytosine also differs to the unmethylated cytosine in the occurring change in current, thus allowing a real time methylation sequencing without any prior labelling or modification [8-12].

To evaluate the reliability of our previous *TRPA1* promoter methylation studies [13, 14], we compared the methylation rates obtained via bisulfite and nanopore sequencing. For this purpose, we used the Cas-mediated PCR-free enrichment to target the *TRPA1* promoter region for subsequent MinION nanopore sequencing. This targeted sequencing approach allows to enrich for loci of interest, yielding in high coverage of the desired genomic regions [15], which is necessary to allow a reliable evaluation of methylation rates. The absolute minimum number of reads required might depend on the target region and the methylation level.

## 2. Results and Discussion

### 2.1. Comparison of the bisulfite and nanopore sequencing methods for methylation calling by repeated measurements in one DNA sample

The comparison between the bisulfite and nanopore sequencing methods revealed similar methylation rates for the seven CpGs of the *TRPA1* promoter region analyzed in repeated measurements of DNA extracted from buffy coat of a healthy volunteer. As shown in figure 1, the mean methylation rates, as well as the methylation rates of the individual replicates (10 for Sanger bisulfite sequencing, 12 for nanopore sequencing) were congruent between bisulfite and nanopore sequencing, although the number of reads varied. The latter is discussed in the last section “Accuracy of methylation data determined via nanopore sequencing in relation to the number of calls per site”.

**Figure 1.**
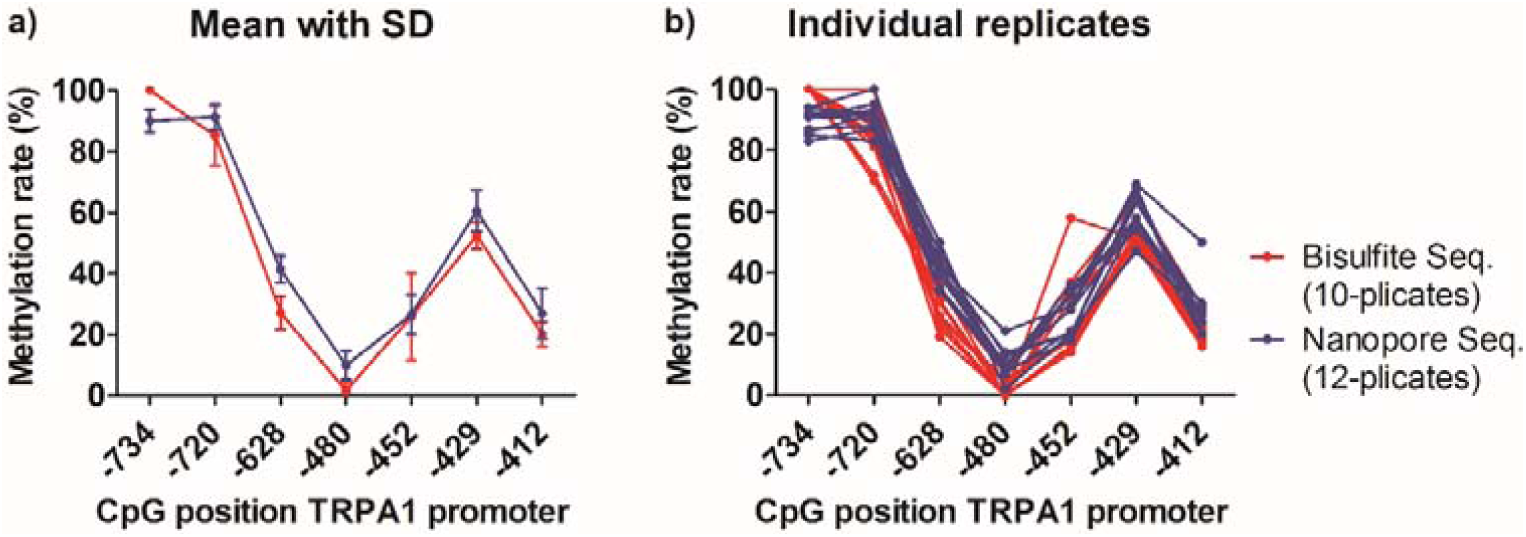
Comparison of methylation rates obtained via direct bisulfite and nanopore sequencing of the *TRPA1* promoter. Sanger bisulfite sequencing was performed in 10-plicates, nanopore sequencing in 12-plicates. a) Mean methylation rates and standard deviation per CpG position, b) Methylation rates per CpG position of single measurements.

Thus, we were able to validate our previous Sanger bisulfite sequencing results of this particular region of the *TRPA1* promoter. Although a variety of new methods for methylation analysis is available, bisulfite sequencing remains a satisfactory and reliable method with single CpG resolution. A study, which compared the performance of widely used methods for DNA methylation analysis that are compatible with routine clinical use in 18 laboratories in seven different countries, found clonal bisulfite sequencing to perform reasonably well, although it did not reach the accuracy and reproducibility of the top-ranking assays analyzed [18]. These assays involved bisulfite conversion, PCR amplification, mass spectrometric quantification, microarray analysis, qPCR with a methylation specific probe, high resolution melting analysis, high-performance liquid chromatography, enzyme-linked immunosorbent assay, and cloning. The authors mention that until recently the method of clonal bisulfite sequencing was considered the gold standard for locus-specific DNA methylation mapping, but suggest using one of the novel, less labor-intensive assays for biomarker development. Amplicon bisulfite sequencing and bisulfite pyrosequencing showed the best all-round performance in this study [18]. Direct bisulfite sequencing is much less labor-intensive than involving plasmid cloning, but unfortunately is accompanied by a decrease of accuracy. The extent of decrease mainly depends on the tools used to calculate the proportion of peak heights between cytosine and thymine. In general, in our study lower levels were measured with bisulfite sequencing in comparison to nanopore sequencing. Differences are most obvious at CpG positions -628, -480 and -429, but are also visible at CpG -720 and -412 (Fig. 1). The only CpG site with higher levels for Sanger bisulfite sequencing compared to nanopore sequencing is CpG -734, which might be due to the inaccurate reading at the beginning of the sequence. The calculated standard deviation of our measurements is below 10 except at CpG -452 (Table 1), which mainly resulted from one outlier of the bisulfite sequencing 10-plicates at this CpG position (Fig. 1). At CpG position -628 the standard deviation is low for both methods. The observation that cytosine modifications seem to have a protective effect against bisulfite treatment-induced DNA degradation, and that bisulfite treatment leads to a depletion of genomic regions enriched for unmethylated cytosines [7], did therefore not concur with our results. A possible explanation for our observation could be a biased amplification of the bisulfite-treated DNA, which is sequence-dependent and often strand-specific [19]. Since the GC-content of methylated DNA after bisulfite treatment is higher than that of unmethylated DNA, an inaccurate estimate of methylation is possible. The higher melting temperature of the DNA with higher GC-content may result in an increased likelihood for secondary structure formation for some sequences, and therefore decrease the PCR efficiency compared to unmethylated sequences [19]. An additional explanation for the lower methylation rates observed with bisulfite sequencing compared to nanopore sequencing could be an inappropriate bisulfite conversion. False negative data occur with longer incubation times leading to higher degradation and accumulation of inappropriate conversion, without necessarily contributing to overall conversion efficiencies [2, 4].

**Table 1.** Mean methylation rates and standard deviation of Sanger bisulfite and nanopore sequencing.

| CpG position | Methylation rate (%) |  |  |  |
| --- | --- | --- | --- | --- |
|  | Bisulfite sequencing<br>(10-plicates) |  | Nanopore sequencing<br>(12-plicates) |  |
|  | Mean | SD | Mean | SD |
| -734 | 100.0 | 0.0 | 90.0 | 3.8 |
| -720 | 85.2 | 9.9 | 91.3 | 4.2 |
| -628 | 27.1 | 5.4 | 41.4 | 4.5 |
| -480 | 1.6 | 2.7 | 10.0 | 4.6 |
| -452 | 25.9 | 14.3 | 26.6 | 6.4 |
| -429 | 52.4 | 4.5 | 60.6 | 6.7 |
| -412 | 20.1 | 4.0 | 26.9 | 8.3 |

### 2.2. Comparison of methylation data obtained via nanopore sequencing with previously published bisulfite sequencing results

In order to verify our previously published Sanger bisulfite sequencing data of the *TRPA1* promoter [13, 14], we measured the methylation rates via nanopore sequencing in some samples of the same cohorts used before. Whereas it was possible to re-extract DNA from healthy control whole blood samples [13], only DNA from Crohn patients extracted previously [14] was available. Only five of the Crohn patient DNA samples possessed a sufficient DNA concentration for nanopore sequencing, but the quality was lower than of the DNA extracted from buffy coat or extracted freshly from whole blood. The lower DNA-integrity of the Crohn patient group (n = 5) gave rise to a relatively low number of calls per site of the seven CpGs analyzed between 16 and 37, whereat the number of calls per site at CpG -628 was between 20 and 30. DNA of the healthy control group (n = 10) gave rise to the number of calls per site for the seven CpGs between 60 and 223, and for CpG -628 between 70 and 199. Although the experimental conditions were therefore not ideal, the methylation rates measured with the two different methods are congruent (Fig. 2). Again, lower methylation levels were obtained with Sanger bisulfite sequencing compared to nanopore sequencing. As discussed above, the reason for this observation might be a biased amplification of bisulfite-treated DNA (compare Fig. 1). According to the findings obtained from the test DNA (Fig. 1, section 2.1), the only CpG site with higher levels for bisulfite sequencing compared to nanopore sequencing is CpG -734, which might be due to the inaccurate reading at the beginning of the sequence. During the quality control of methylation data carried out in our previous studies, CpG sites with less than 5 % inter-individual variability, which applied to CpGs -734 [13] and -480 [13, 14], were rejected. However, the methylation rates of all subjects for CpG -734 were measured around 100 %, and those for CpG - 480 around 0 % (data not shown). Our direct Sanger bisulfite sequencing approach was not applied for diagnostic purposes of individuals but to gain an overview about the methylation rates of single CpG sites in relation to the subjects’ pain sensitivities within a large cohort [13, 14]. Since a possible bias would apply to all samples independent of the subjects’ pain sensitivity, minor discrepancies are acceptable.

**Figure 2.**
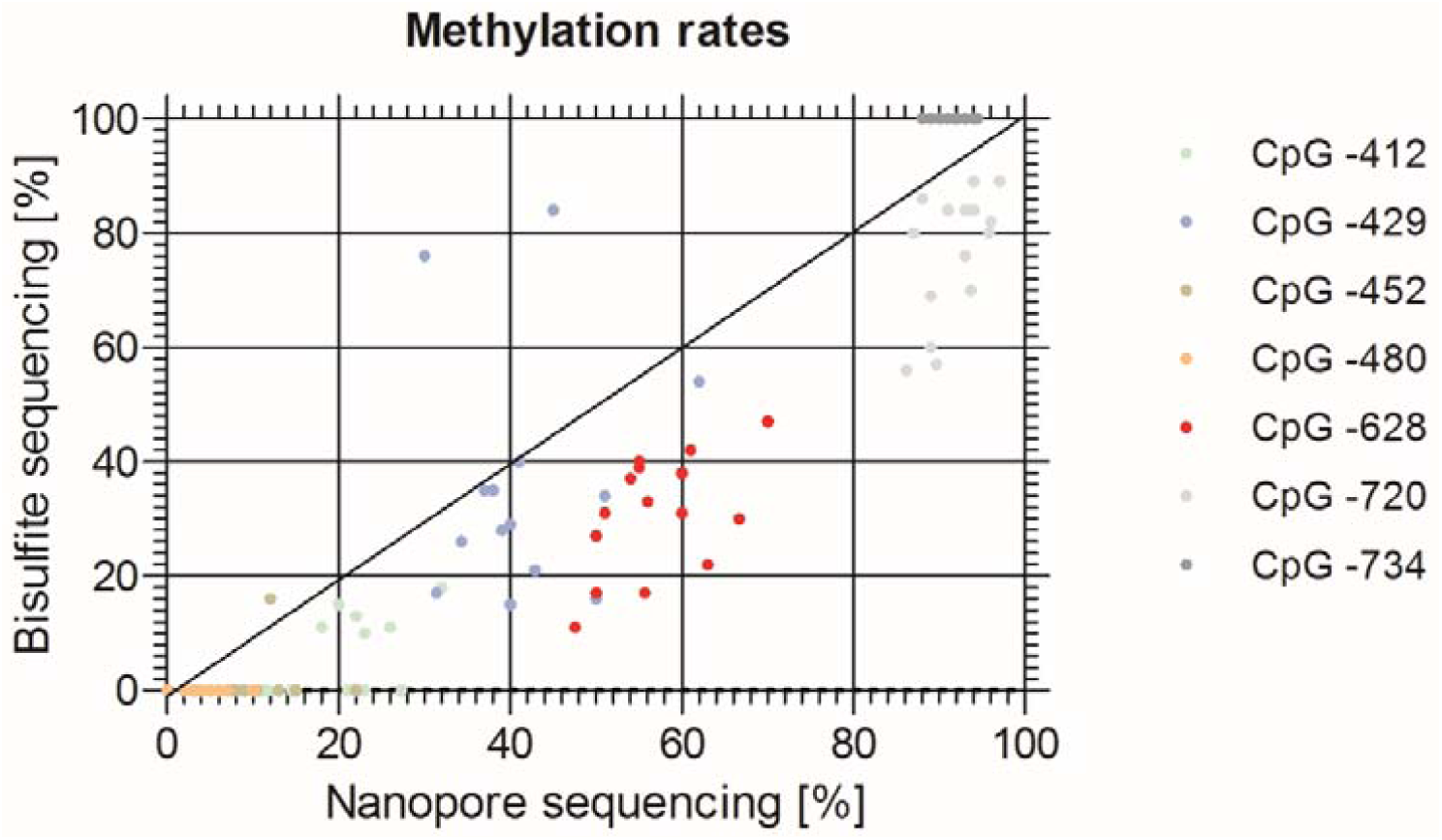
Comparison of methylation data of the *TRPA1* promoter obtained via bisulfite and nanopore sequencing in two cohorts. Sanger bisulfite sequencing results were published previously [13, 14], nanopore sequencing was conducted in the present study.

Figure 3 shows the correlation between CpG -628 methylation and pressure pain threshold, measured previously by an algometer over the thenar muscles [13, 14]. For Sanger bisulfite sequencing data obtained from our previous studies [13, 14], the correlation is more pronounced than for nanopore sequencing. For both methods, it did not reach significance level, which is in contrast to the results from our previous studies [13, 14]. This could be due to the very small cohort size resulting from the limited availability of samples with a sufficient quality for Nanopore sequencing (10 healthy subjects, 5 Crohn patients). However, a trend for low pressure pain thresholds (high pain sensitivities) with high methylation rates at CpG -628 is still visible.

**Figure 3.**
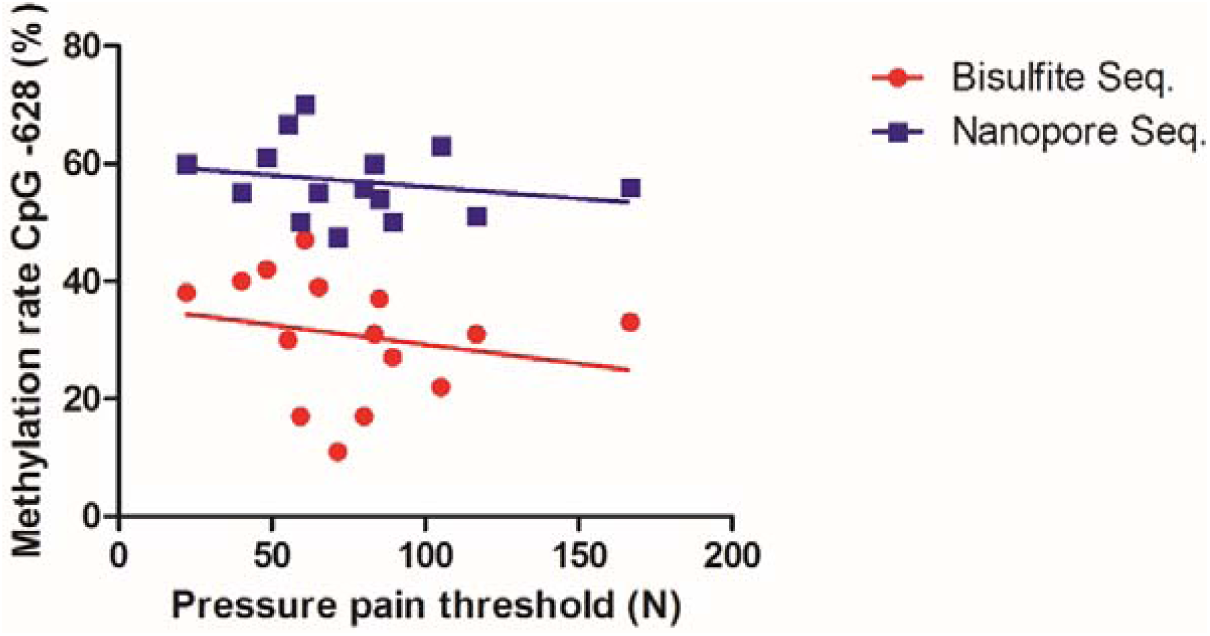
Correlation between CpG -628 methylation rate, determined via bisulfite and nanopore sequencing, and pressure pain threshold. Sanger bisulfite sequencing results and pressure pain thresholds were published previously [13, 14], nanopore sequencing was conducted in the present study. Bisulfite sequencing: R^2^ = 0.049, p = 0.426; nanopore sequencing: R^2^ = 0.048, p = 0.435.

### 2.3. Accuracy of methylation data determined via nanopore sequencing in relation to the number of calls per site

Whereas for DNA sequencing irrespective of base modification calling, obtaining a high number of calls per site is of less importance, a sufficient number is necessary to assess the accurate percentage values of methylation in a mixture of DNA strands. Although the usage of several guide RNAs upstream and downstream of the target region is recommended [15], surprisingly, a single pair of guide RNAs (# 4 and # 9) was shown to result in the highest number of calls per site obtained. An overview of the relative positions of tested guide RNAs to the target region within the *TRPA1* promoter is shown in figure 6a. In addition to the guide RNA strategy, the number of calls per site highly depends on the concentration and the quality of the DNA used for nanopore sequencing. Therefore, as mentioned before, the number of calls per site obtained when measuring the remaining DNA from Crohn patients extracted for our previous study [14], was relatively low. However, during establishment of the most suitable sgRNA strategy for our region of interest, we first combined all of the guide RNAs designed (Fig. 4a and table 2), thereby generating a lower number of reads as compared to the sole application of guide RNAs # 4 and # 9, using the same DNA control sample extracted from buffy coat. Usage of guide RNAs # 1, # 3 and # 8 probably hampers adapter binding, and thereby decreases the efficacy of the following sequencing reaction due to persisting Cas9 molecules within the region to be read. Despite a thermal Cas9-deactivating step in the protocol, the Cas9 enzyme, being known to hardly dissociate from the DNA after cutting [20], remains bound to the DNA strand [15]. Since persistent Cas9 binding was shown to block DNA repair proteins from accessing Cas9-generated breaks [21], this observation might be valid also for the experimental conditions in Cas9-mediated PCR-free enrichment protocols. In fact, it has already been described that adapters bind preferentially on the 3’-side of a Cas9 cut, as the enzyme remains stably bound to the 5’-side of the sgRNA [15]. It is conceivable that the first subsequent step after cutting, namely dA-tailing, is hindered due to persisting Cas9 molecules, since the polymerase requires at least 4 bp to bind to a DNA strand. Moreover, the DNA is single-stranded within the Cas9 molecule, and one of the strands is even hybridised with the guide RNA. Therefore, the prerequisites for polymerase-binding are not fulfilled. Since y-“Sequencing adapters are ligated primarily to Cas9 cut sides, which are both 3’ dA-tailed and 5’ phosphorylated” (manual “Cas9 targeted native barcoding”, Oxford Nanopore Technologies) [22], adapter ligation to DNA strands with bound Cas9 will be less effective. In addition, the phosphate group is also most likely inaccessible with persisting Cas9. All these factors make a reduced sequencing capability comprehensible for persistent Cas9 molecules. Since Cas9 binding occurs at the 5’-side of the guide RNA, and cutting close to the PAM region at the 3’-side, the blocking of adapter binding due to persisting Cas9 molecules after cutting is locatable in relation to the target (compare Fig. 4b and c). This should be considered when choosing sgRNAs for using the Oxford Nanopore Technologies “Cas-mediated PCR-free enrichment”-protocol. As depicted in figure 4b, the usage of sgRNAs # 4 and # 9 results in accessibility of both ends for adapter ligation after cutting, which is in contrast to the situation depicted in figure 4c. Therefore, the addition of sgRNAs # 1, # 3 and # 8 leads to a reduced number of reads, since according to statistical probability some DNA molecules will be blocked (at least unilaterally) for adapter ligation.

**Figure 4.**
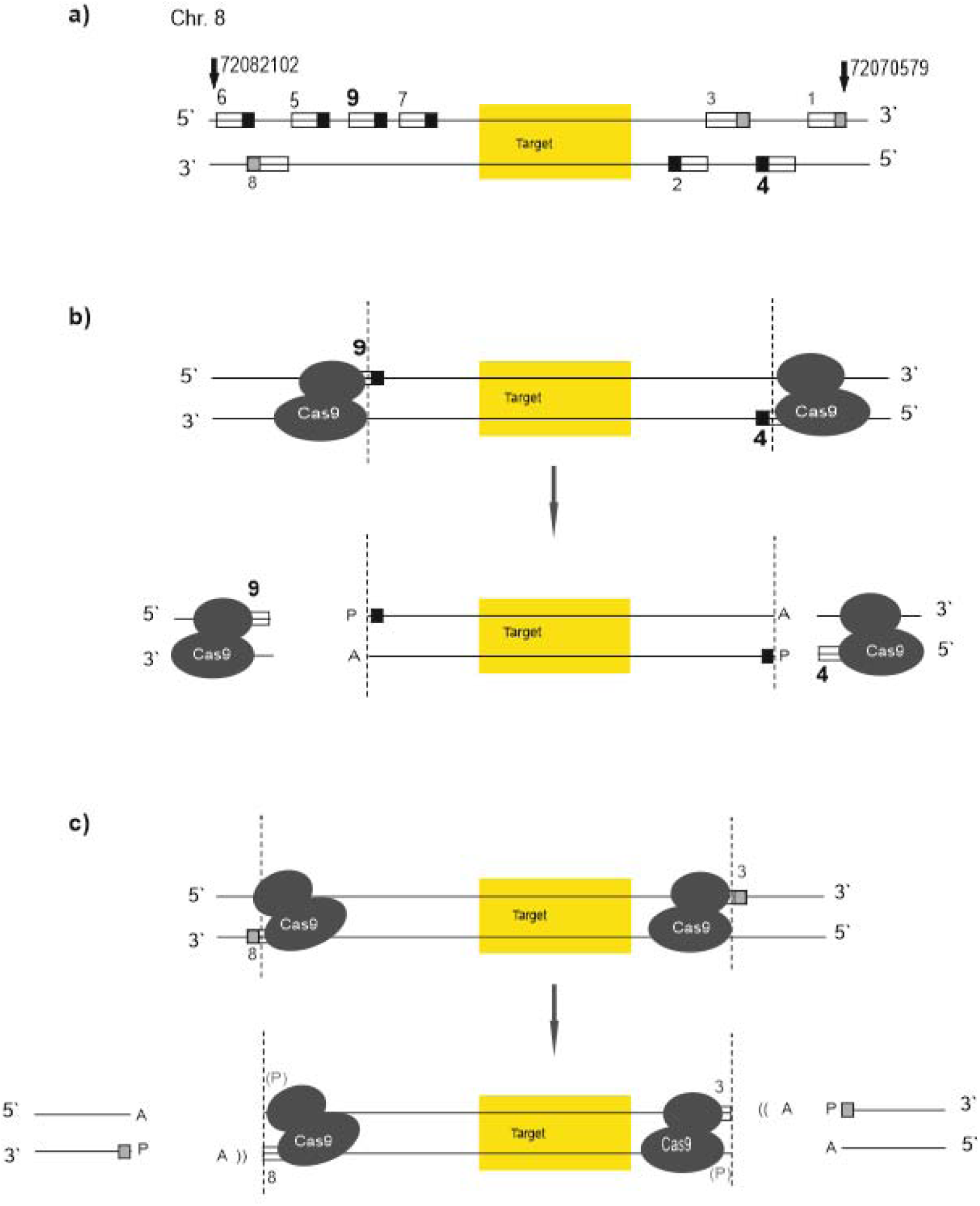
Positions of guide RNAs designed for the target *TRPA1* promoter. Guide RNAs are numbered (compare table 3) and are depicted as boxes, PAM motifs are indicated by black squares, PAM motifs of “interfering” guide RNAs are indicated by grey squares. The guide RNAs yielding the highest number of calls per site (# 4 and # 9) are printed in bold. a) Position of guide RNAs relative to the target region on chromosome 8. GRCh38 positions are marked by arrows. b) Correct position of guide RNAs enabling adapter ligation after cutting. c) Blocking of adapter binding by suboptimal positioned guide RNAs resulting in residual Cas9 molecules within the target region.

**Table 2.**
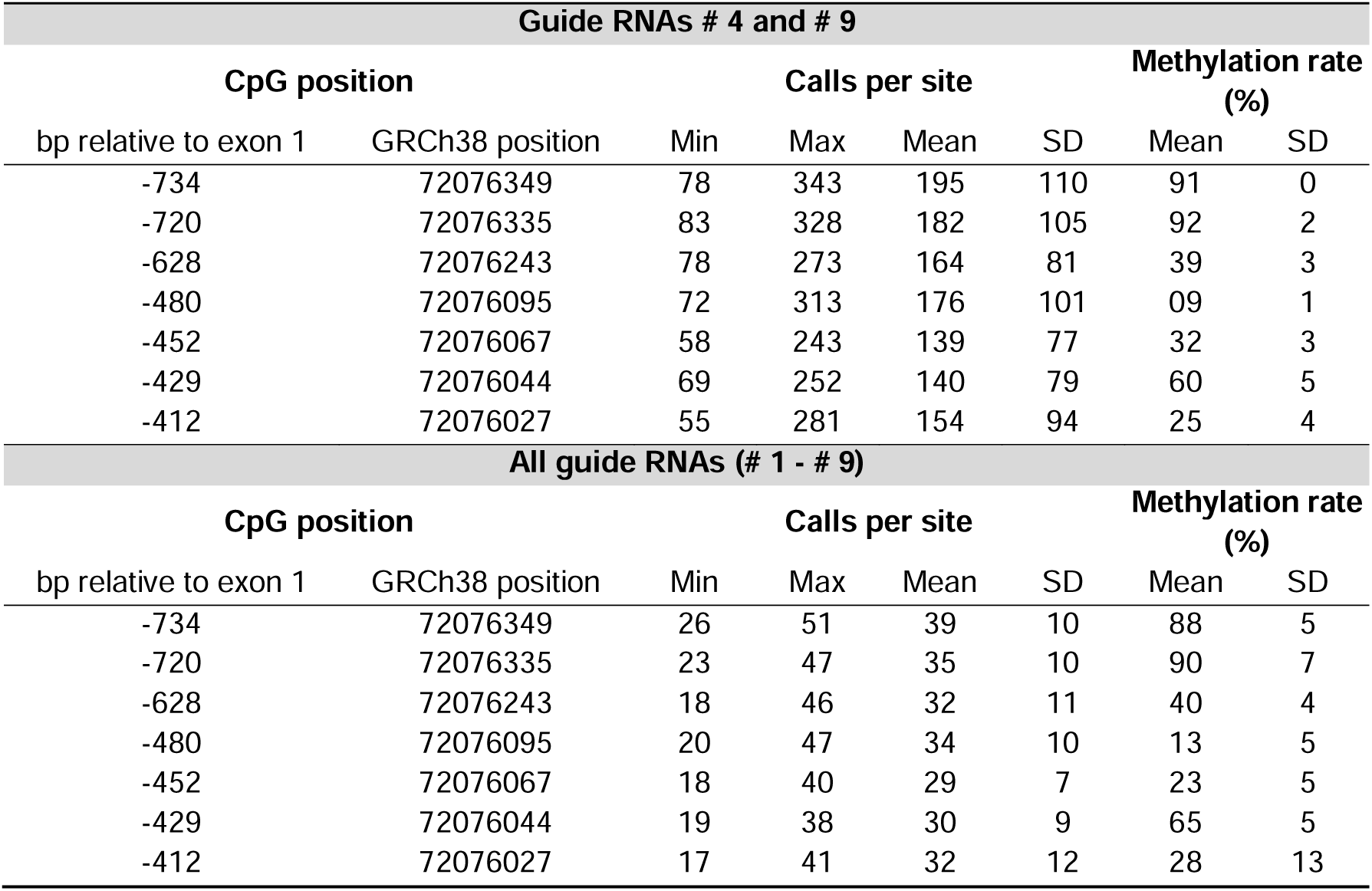
Calls per site and methylation rates obtained applying two different guide RNA strategies. Data with guide RNAs # 4 and # 9 are based on four experiments (number of calls per site 55-343), with all guide RNAs on five experiments (number of calls per site 17-51). The same DNA sample extracted from buffy coat was used for all sequencing experiments.

Despite the lower numbers of calls per site obtained by utilization of the “interfering” guide RNAs # 1, # 3 and # 8, methylation rates were measured within the same range for both guide RNA strategies (Table 2). Thus, nanopore data of the Crohn patient DNA with poor integrity can be assumed to be reliable, despite lower read numbers.

Nevertheless, the representation of data points in figure 5 reveals that the measurement of methylation rates is not completely stable before reaching a number of calls per site of 100 (or a bit less) for all of the CpGs analyzed. However, deviations are not high, even in the case of low read numbers. According to Oxford Nanopore Technologies a read depth of around 30 x is advised for phasing methylation calls. Apart from the influence of DNA concentration and integrity, as well as the chosen guide RNAs, also the condition of the flow cell and the corresponding number of active pores has an impact on read numbers. Thus, variations in the number of calls per site may occur regularly. Since one flow cell with a maximum load of five DNA samples can be used per nanopore run, it is not possible to circumvent the resulting divergence.

**Figure 5.**
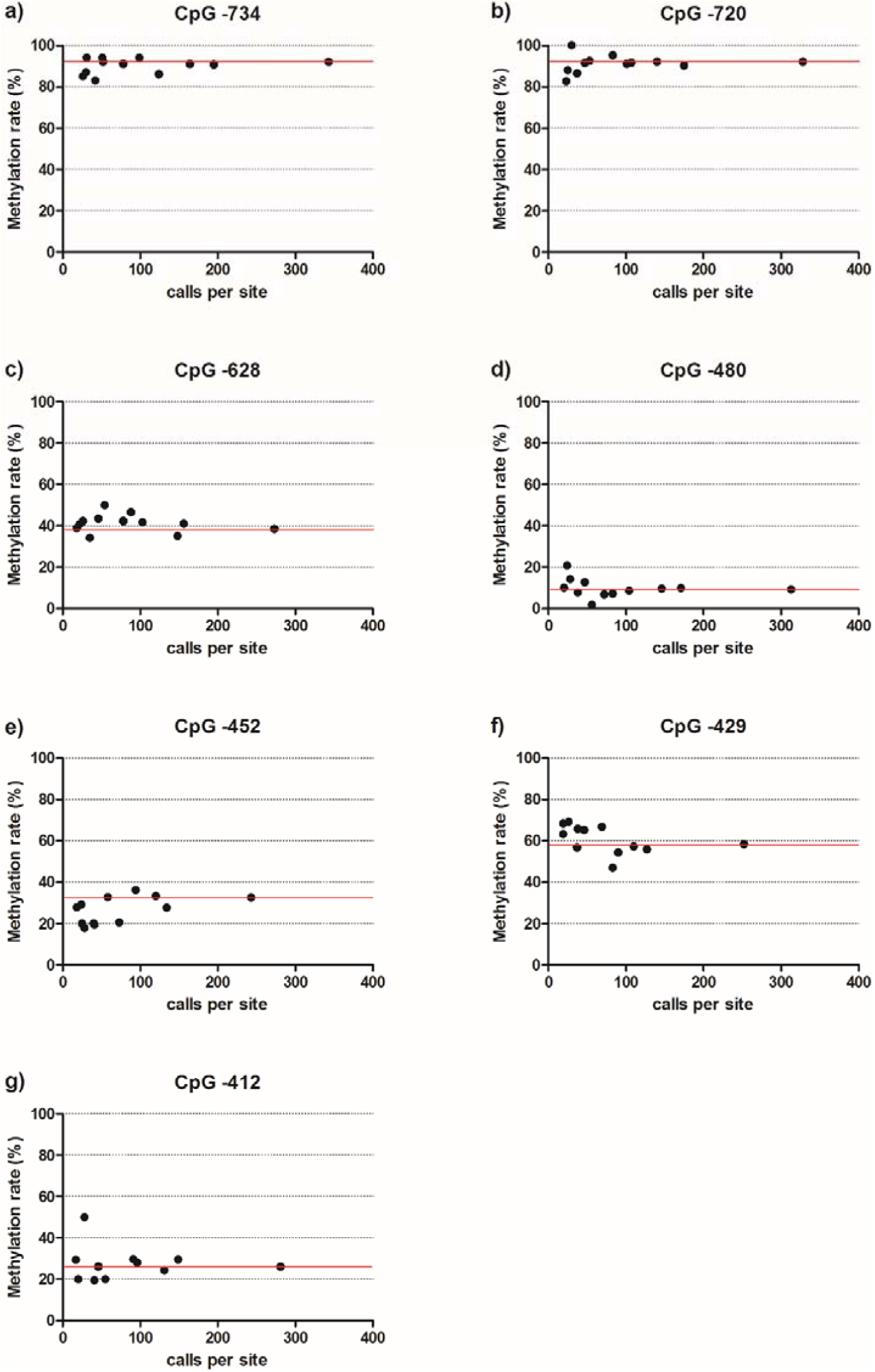
Methylation rates of the *TRPA1* promoter determined via nanopore sequencing in relation to the number of calls per site. Data based on 12 measurements of the same DNA sample. The red line marks the methylation rate measured with the highest number of calls per site per CpG, which is presumably the most accurate value. Different flow cells and different guide RNA strategies for Cas-mediated PCR-free enrichment were used leading to varying numbers of calls per site.

To conclude, according to the nanopore sequencing results, we obtained reliable results with the chosen method of direct Sanger bisulfite sequencing in our previous studies [13, 14]. The method of direct bisulfite sequencing is suitable to gain an overview of the underlying regulatory interrelation when analyzing large cohorts, although for diagnostic purposes other methods should be used to allow valid statements for an individual. The advantages and disadvantages of the method of choice should be evaluated carefully before starting the analyses. Whereas direct Sanger bisulfite sequencing enables a high throughput of samples (2 × 96) when analyzing a single promoter region, MinION nanopore sequencing is labor- and time-intensive due to the maximum load of five samples per flow cell. However, utilization of sequencing devices, which are able to run several flow cells at once, is possible. Methylation data obtained via nanopore sequencing are more accurate compared to those obtained via direct Sanger bisulfite sequencing (at least when reaching sufficient numbers of reads), due to the absence of bisulfite conversion and PCR amplification. Nanopore sequencing is more suitable when analyzing the promoters of a panel of several genes within each sample, rather than analyzing one single promoter region. However, methylation data obtained via direct Sanger bisulfite sequencing are adequate to draw conclusions in large cohort studies.

## 3. Materials and Methods

### 3.1. Samples

Blood for comparison of methylation rates obtained via Sanger bisulfite and nanopore sequencing in 10-plicates and 12-plicates, respectively, was drawn from a healthy volunteer recruited in Hannover prior to DNA extraction from buffy coat by the Hannover Unified Biobank. Approval for analysis of the healthy control was obtained at the Ethics Committee of the Hannover Medical School (Permit Number 2842-2015).

For comparison with Sanger bisulfite sequencing results of healthy subjects from a previous study [13], DNA extraction from whole blood was performed as stated below (n = 10) in order to obtain DNA of high quality and yield for nanopore sequencing. For comparison with Sanger bisulfite sequencing results of Crohn patients from another previous study [14], the DNA that was already extracted from whole blood as stated in the published manuscript was used (n = 5), due to the unavailability of blood samples.

### 3.2. DNA extraction

800 µl of whole blood were incubated with 80 µl Proteinase K (Macherey-Nagel, Düren, Germany) for 15 min at room temperature and centrifuged at 21 000 xg and 4°C for 5 min. The resulting 200 µl-fractions were separated, and the Nucleo-Mag Blood 200 µl DNA Kit (Macherey-Nagel, Düren, Germany) was used for extraction and clean-up of genomic DNA. For pipetting and transferring steps, as well as for purification of DNA a Biomek N x P (Beckman Coulter, Brea, CA) was used. DNA concentration was determined on a DeNovix DS-11 Spectrophotometer (DeNovix, Wilmington, USA), and the fraction with the highest DNA concentration of each sample was used for nanopore sequencing.

### 3.3. Sanger bisulfite sequencing

DNA samples were bisulfite converted and purified using the EpiTect 96 Bisulfite Kit (QIAGEN, Hilden, Germany). Amplification of the *TRPA1* promoter target sequence using the forward primer 5’-GTTTGTATTAGATAGTTTTTTTGTTTG-3’ and the reverse primer 5’-TCCTACAAACCTATATTTCCCAC-3’, purification of the amplified target sequence and sequencing on a 3500xL Genetic Analyzer (ABI Life Technologies, Carlsbad, USA) was performed as described previously [13]. All samples showed a quality value above 20 for trace score in the Sequence Scanner Software (ABI Life Technologies, Carlsbad, USA). Methylation rates for each CpG site were determined via the Epigenetic Sequencing Methylation Analysis Software [16].

### 3.4. Nanopore sequencing

Guide RNAs (Table 3) were designed using the target prediction program chopchop (http://chopchop.cbu.uib.no) and ordered from IDT (Integrated DNA Technologies, Coralville, USA).

**Table 3.**
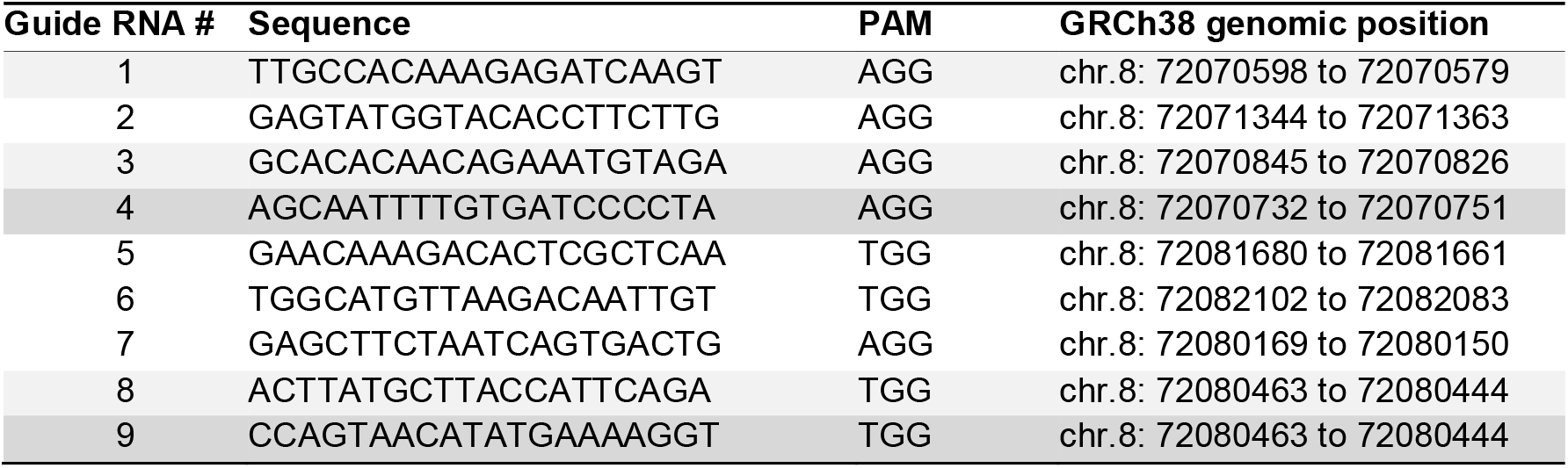
Guide RNAs used for establishing the Cas-mediated PCR-free enrichment of the *TRPA1* promoter sequence analyzed via nanopore sequencing. Different combinations were tested, with a combination of guide RNA # 4 and # 9 giving the highest number of calls per site. Guide RNAs # 1, # 3 and # 8 were used as “interfering” guide RNAs as a control. PAM: Protospacer Adjacent Motif.

For the selection of guide RNAs the following quality criteria were applied: efficiency ≥ 0.5, self-complementarity max. 2, GC content 40-70 %, mismatches between off-targets and guide RNA: MM0 (no mismatch) = 0, MM1 (1 mismatch) = 0, MM2 (2 mismatches) as low as possible, MM3 (3 mismatches) as low as possible. Several guide RNAs per cutting site were identified and subsequently the quality was assessed using the online tool Off-spotter (https://cm.jefferson.edu/Off-Spotter). Guide RNAs with off-target mismatches close to the PAM (protospacer adjacent motif) were preferred over those with off-target mismatches far from the PAM due to a reduced binding and cutting probability of the Cas9 enzyme. The DNA quality was assessed via pulsed-field gel analysis using a Pippin Pulse electrophoresis power supply (Sage Science, Beverly, USA). 5 µg of high-molecular weight DNA of each sample were used for the Cas-mediated PCR-free enrichment using the Ligation Sequencing (SQK-LSK109) Kit (Oxford Nanopore Technologies, Oxford, UK) and the Native Barcoding Expansion 1-12 (PCR-free) (EXP-NBD104) Kit (Oxford Nanopore Technologies, Oxford, UK) in the following order: Dephosphorylating genomic DNA, Preparing the Cas9 ribonucleoprotein complexes (RNPs), Cleaving and dA-tailing target DNA, Native barcode ligation, Adapter ligation (during this step buffer AMII instead of buffer AMX and no nuclease-free water was added to the ligation mixture due to prior barcoding), AMPure XP bead purification (TE buffer and AMPure XP beads were scaled up due to the higher final volume after barcoding; the pellet was resuspended in 14 µl of preheated elution buffer at 37°C for 20 minutes with flicking the tube every 5 minutes), Priming and loading the SpotON flow cell FLO-MIN106D (Oxford Nanopore Technologies, Oxford, UK), and starting the sequencing run on the MinION (Oxford Nanopore Technologies, Oxford, UK). Raw sequencing data was basecalled using guppy v. 2.7 (Oxford Nanopore Technologies, Oxford, UK) with standard settings and config file ‘dna_r9.4.1_450bps_hac’. Basecalled reads were aligned to GRCh38 using minimap2 [17]. Methylation calling and determination of methylation frequency was performed using nanopolish v 0.12.4 [11].

### 3.5. Statistical analyses

For statistical calculations and data illustration Prism 5 (GraphPad) was used. For correlation analysis of methylation data between bisulfite and nanopore sequencing, as well as between CpG -628 methylation and pressure pain threshold, a p value of ≤ 0.05 was considered significant.

## Author Contributions

Conceptualization, H.F. and A.L.; methodology, K.J., H.P., A.B., S.G, G.S., L.W., B.B. and F.J.M.; software, M.D., G.S. and C.D.; validation, S.G., K.J., H.P. and A.B.; formal analysis, S.G., K.J. and M.D.; investigation, K.J. and S.G.; resources, H.F., L.W. and F.J.M.; writing – original draft preparation, S.G. and K.J.; writing – review and editing, A.B., H.P., H.F., M.D., G.S., L.W., C.D., B.B., F.J.M. and A.L.; visualization, S.G. and K.J.; supervision, H.F. All authors have read and agreed to the published version of the manuscript.

## Institutional Review Board Statement

The study was conducted according to the guidelines of the Declaration of Helsinki, and approved by the Ethics Committee of the Hannover Medical School (Permit Number 2842-2015).

## Informed Consent Statement

Informed consent was obtained from all subjects involved in the study.

## Acknowledgments

We would like to acknowledge the assistance of the Hannover Unified Biobank for DNA extraction from buffy coat, and thank Roland Meister (Institute of Experimental Ophthalmology, University Eye Hospital, Hannover Medical School, Hannover, Germany) for technical support. We would like to thank Tino Münster (Department of Anesthesiology, Friedrich-Alexander University Erlangen-Nuremberg, Erlangen, Germany; Clinic for Anesthesiology and Critical Care, Hospital of the Order of St.John of God Regensburg, Regensburg, Germany) and Andreas Winterpacht (Institute of Human Genetics, Friedrich-Alexander University Erlangen-Nuremberg, Erlangen, Germany) for providing blood samples of healthy individuals and DNA samples of Crohn patients for reanalysis via nanopore sequencing.

